# Control and coding of pupil size by hypothalamic orexin neurons

**DOI:** 10.1101/2022.04.12.488026

**Authors:** Nikola Grujic, Alexander Tesmer, Ed F. Bracey, Daria Peleg-Raibstein, Denis Burdakov

## Abstract

Brain orexin (hypocretin) neurons are implicated in sleep-wake switching and reward-seeking, but their roles in rapid arousal dynamics and reward perception remain unclear. Here, cellspecific stimulation, deletion, and in vivo recordings revealed strong correlative and causal links between pupil dilation - a quantitative arousal marker - and orexin cell activity. Coding of arousal and reward was distributed across orexin cells, indicating that they specialize in rapid, multiplexed communication of momentary arousal and reward states.

---

The orexin (hypocretin) system of the lateral hypothalamus (LH) projects brain-wide, with particularly strong connections to arousal and reward centres (1–3). Through these connections, the orexin network regulates sleep-wake switching and autonomic function, as well as feeding and exploratory behaviours (4–9). Pupil dilation is routinely used in human experiments as a measure of arousal and autonomic function (10), and for predicting key aspects of cognition, such as the exploration-exploitation balance (11, 12). As such, and with its strong tracking of the locus coeruleus (LC) noradrenergic system (13, 14), pupil size measurements have been used to support the adaptive gain theory of arousal (12, 15). Pupil dilation has also been correlated to the activity of cholinergic (14) and serotonergic (16) neuro-modulatory systems. However, orexinergic modulation of moment-to-moment arousal, as indexed by pupil dilation, has not been explored. In particular, the related question of interplay of arousal and reward representations in individual orexin neurons (9) has not been answered experimentally.

To causally test the pupil dilation responses elicited by orexin cell activation, we selectively photo-stimulated LH orexin neurons, while tracking pupil diameter in anaesthetized mice (Fig. 1a, Methods). We observed rapid dilation in response to stimulation, which declined after stimulation offset (Fig. 1b-d, statistics are given in the figure legends). The effect was stimulation-frequency dependent, with higher frequency stimulation eliciting faster and greater pupil dilation (Fig. 1b-d).

**Fig. 1:**
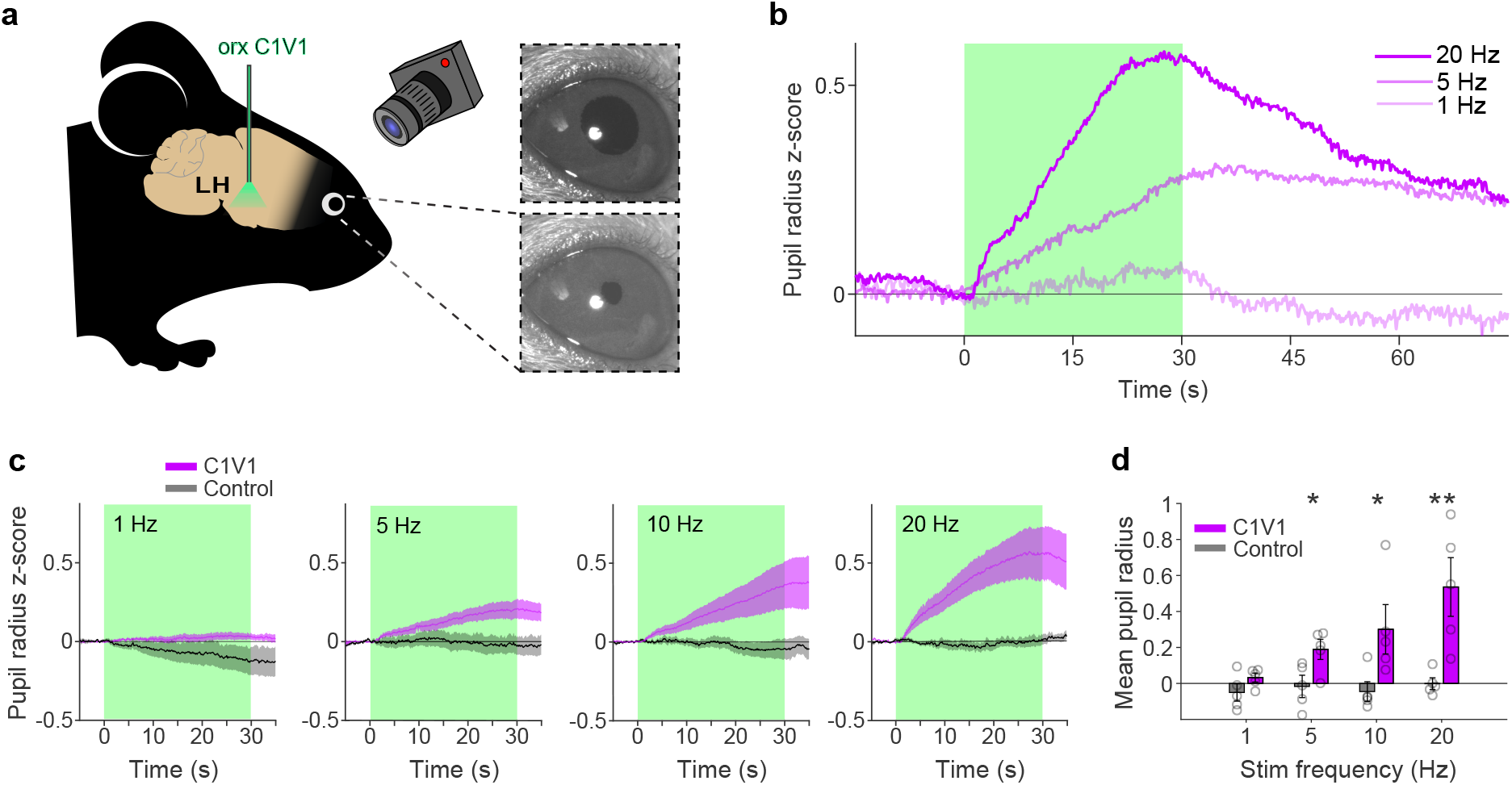
Orexin cell stimulation elicits pupil dilation. **a)** Mouse pupils were recorded under isoflurane anesthesia during optogenetic photo-stimulation of orexin neurons in the LH. **b)** Example pupil responses of one mouse to different stimulating frequencies. Duration of stimulation is indicated by the green box. **c)** Pupil responses for the opsin (n = 5 mice) and control (n = 5 mice) groups at increasing stimulation frequencies (left to right, shaded areas show SE). **d)** Mean pupil sizes for the last 10 seconds of stimulation, asterisks indicate significant differences between groups (two-tailed t-test, left to right frequencies: t = 1.6, 2.7, 2.6, 3.6; p = 0.15, 0.02, 0.03, 0.006).

Next, we tested how disruption of orexin neuropeptide signalling affected pupil size in awake and anesthetized animals. To distinguish the contribution of orexin neurotransmission to pupil dilation from other neurotransmitters emitted by orexin neurons, we repeated the optogenetic stimulation experiment (Fig. 1) while specifically blocking orexin receptors with the antagonist Almorexant (ALM, Fig. 2a, Methods). ALM reduced both tonic pupil dilation (Fig. 2b,c), and the extent of pupil dilation during the orexin cell photo-stimulation evoked, phasic response (Fig. 2d). Interestingly, rapid dilation at photo-stimulation onset was initially similar between ALM and vehicle conditions (Fig. 2d), but diverged after 1 - 2 seconds, with slower and smaller dilation occurring in ALM-injected mice thereafter (Supp. Fig. 1). This suggests that fast transmitter(s) released by orexin cells (17) caused the initial dilation. Overall, these results establish a causal link between orexin neurotransmission and pupil size.

**Fig. 2:**
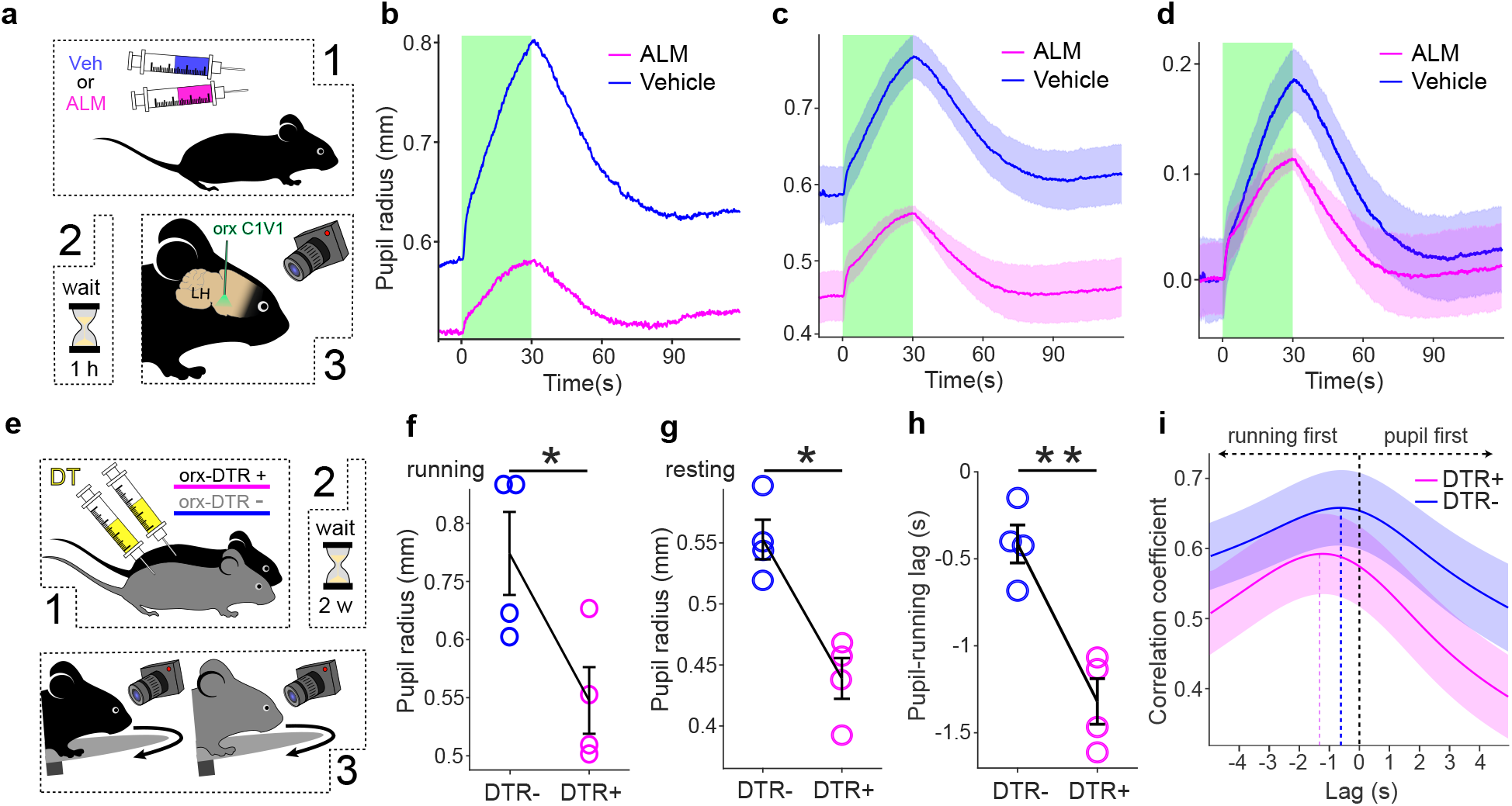
Disruption of orexin cell function leads to observable pupil size differences. **a)** The pupil was recorded during 20 Hz photo-stimulation of LH orexin neurons in Almorexant (ALM) or vehicle injected, isoflurane anaesthetized mice. **b)** An example pupil size trace from one mouse during orexin cell photo-stimulation after ALM or vehicle injection. Green shaded area indicates photostimulation. **c)** Mean absolute pupil size across mice (n = 4 mice). SEs are shown by shaded areas. **d)** Mean pupil dilation across mice, with baseline subtracted. **e)** Pupil radius of orx-DTR+ (n = 4 mice) and orx-DTR- (n = 4 mice), DT injected mice was recorded during running on a wheel. **f,g)** Mean pupil size during running (left, one-tailed t-test, t = 2.3, p = 0.03) and no-running epochs (right, one-tailed t-test, t = 4.8, p < 0.01). Error bars show SE. **h)** Mean pupil radius to running onset lag at maximum correlation values for the two groups. Error-bars show SE (one-tailed t-test, t = 5.6, p < 0.01). **i)** Cross correlation analysis of pupil radius and associated running signals. Mean correlation coefficients at different lags for the two groups. SEs are shown by shaded areas.

To investigate the effect of orexin neurons themselves on pupil modulation, we selectively ablated them using the orx-DTR mouse model (see Methods). We imaged pupils of orexin cell ablated, head-fixed mice, allowed to run freely on a wheel (Fig. 2e). Due to the previously established link between orexin cell activity and locomotion (7), we investigated the effect of orexin cell deletion on the relationship between pupil size and spontaneous locomotion. We found pupil size was significantly reduced in orexin-cell-ablated mice both during running and resting (Fig. 2f, g). Furthermore, in a cross-correlation analysis, the lag from running to pupil dilation was significantly increased in orexin-cell-ablated mice (Fig. 2h,i). This result shows that orexin neurons are necessary for normal control of pupil size in behaving animals.

Orexin cell activity co-varies with multiple factors, including reward and locomotion (7, 9). To investigate pupil size coding in individual orexin cells in relation to these variables in behaving mice, we used volumetric 2-photon GRIN lens imaging in LH. We monitored orexin cell-targeted GCaMP6s activity, while the mouse was freely running on a wheel and receiving milkshake rewards at random intervals (Fig. 3a). We found that pupil size followed orexin cell activity closely, with cell activation preceding dilation (Fig. 3b,c, Supp. Fig. 2a,c). Our results revealed cells positively or negatively correlated with pupil size (pupil ON and pupil OFF cells, Fig. 3b). These cell types were pooled for further analysis because no obvious differences were found in the way they coded for the other investigated variables (Supp. Fig. 3.). We also found cells whose activation was associated with reward consumption (Supp. Fig. 4). To quantify and compare the contribution of pupil size, running speed and reward consumption (“predictors”) to individual orexin cell responses, we used an encoding model based on multivariate linear regressions (18) (Fig. 3d). By removing each of the predictors from the encoding model for each cell, and comparing the resulting r^2^ with the complete model, we could infer contributions of each predictor to the explained variance of that cell’s activity (Fig. 3f). This revealed distributed coding of pupil size and reward across orexin neurons. Some cells coded for both pupil and reward, but other cells represented only one of these variables, with a larger proportion of cells coding purely pupil with little reward contribution (Fig. 3e). This is in line with the predictions about the dichotomy of orexin function; representing both arousal and reward (9). In addition, we saw that, within the investigated variables, pupil size contributed most to explained variance in cell activity (Fig. 3g). These results establish pupil size as a strong readout of orexin activity, and display how rapid representations of arousal, reward, and movement are distributed across orexin neurons.

**Fig. 3:**
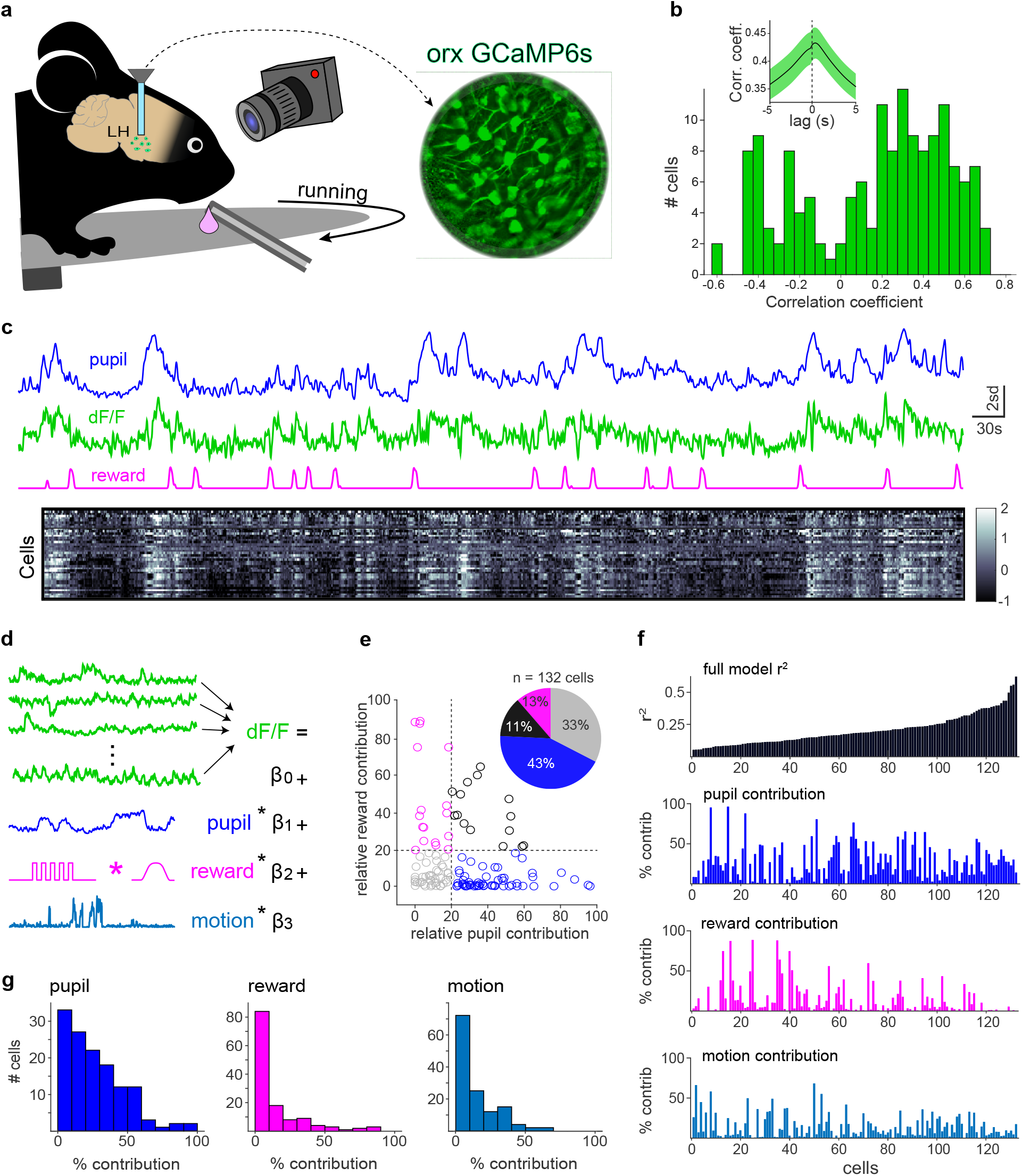
Coding of pupil size and other variables in individual orexin cells. **a)** Pupil is recorded during orx-GCaMP6s, 2-photon, GRIN lens imaging of orexin neurons (n = 150 cells from 3 mice) during free running and reward consumption. **b)** The distribution of cell activity correlations with pupil size (n = 150 cells; the inset shows cross-correlation of pupil ON cells from c, n = 42 cells). **c)** Time aligned activity traces from one mouse. From top to bottom: pupil size, orexin neuron activity, convolved licking trace and a heatmap of activity of all ON-type orexin neurons in the session. **d)** The encoding model. Linear regression was used to quantify the linear relationships between measured variables and activity of each cell separately. **e)** Scatterplot of relative contributions of pupil size and reward consumption to explained variance in each cell’s activity (each point is a separate cell). Pie chart shows the distribution with 20% contribution cut-off. **f)** Top to bottom: the explained variance of each cell’s activity in a model encompassing all investigated variables, pupil, reward, and locomotion percent contributions to the explained variance. **g)** Distributions of cell contributions for each of the investigated variables.

To compare our findings in the orexin system with the noradrenergic system, which is more conventionally associated with pupil dilation, we also investigated pupil tracking of either orexin or LC noradrenaline neuron activity using population-level fibre photometry in behaving mice. We found strong correlations between pupil and both LC noradrenaline and orexin neural activity, with no difference in correlation strength (Supp. Fig. 2a,b,c). There was little difference in coherence between LC or orexin activity and pupil size, with strong coherence at frequencies less than 0.1 Hz, similar to results reported for LC noradrenergic neurons (14) (Supp. Fig. 2d).

In summary, we found strong causative and correlative evidence implicating the orexinergic system in the control of pupil dilation. Our data reveal both tonic and phasic effects of orexin cell activity on pupil size. Orexin neurotransmission was important for increasing pupil dilation, and it is likely than an orexin-cell derived, fast transmitter (probably glutamate (17)) was responsible for rapid-onset effects. Deletion of orexin neurons caused the pupil to be more constricted during both running and rest. In addition, the well documented relation between pupil and running (14) was disrupted by orexin cell loss, suggesting an impairment in the regulation of arousal during activity-state changes. In contrast, orexin cell activation promoted rapid pupil dilation. The majority of individual orexin neurons strongly correlated with pupil, and displayed distributed coding of pupil size, reward, and locomotion. These results shed light on the dichotomy between arousal and reward coding in the orexin system (9), showing that both can be represented in the same neurons, with the extent of each representation varying between neurons.

With their brain-wide projecting axons, orexin neurons might cause pupil dilation by potentiating the activity of several downstream areas. Particularly, this effect could be mediated through strong reciprocal connectivity with the LC (19, 20) or, more directly, via visuomotor cell groups in the brainstem, innervated by orexin neurons (21). However, the relative roles of these neuronal populations in the pupil control circuit would need to be investigated in future studies using connectivity mapping and high-temporal resolution electrophysiology, where exact temporal relationships can be disentangled. Since we show similar relations between pupil size and the LC noradrenaline and orexin cell activity, it will be important to investigate whether some functions previously attributed to LC based on pupil measurements could also be attributed to orexin cell activity, such as arousal gain control (12, 15). Orexin neurons could partially mediate increases in arousal gain associated with exploratory behaviour, and the effect of reward consumption in orexin neurons could be key for transitioning along the exploration-exploitation axis. Furthermore, slow metabolic state signalling to orexin neurons via nutrients and hormones (22–24) would place orexin as a centrepiece in mediating these transitions during, for example, foraging behaviours. The implications of the present results may also be useful for diagnosing orexin cell loss, which occurs in human narcolepsy and is currently diagnosed through highly invasive procedures (25), as well as give insight into other neurological disorders (26).

## Methods

### Animals and surgery

All animal experiments were performed in accordance with the Animal Welfare Ordinance (TSchV 455.1) of the Swiss Federal Food Safety and Veterinary Office, and approved by the Zurich Cantonal Veterinary Office. Subjects were n = 28 adult C57BL/6 mice (at least 8 weeks old). Mice were kept on a reversed 12h/12h light-dark cycle, and all experiments were performed during the dark phase. For all surgeries, mice were anesthetized with 5% isoflurane, after which they were transferred to the stereotaxic surgery setup and maintained on 1.5-2% isoflurane. They were given analgesic and lidocaine was applied to the scalp. An incision was made to access the cranium and two small burr holes were made above the lateral hypothalamus (LH) over both hemispheres (0.9 mm lateral and 1.4 mm posterior from bregma). In the case of GRIN lens implants, only one larger (0.5 mm radius) circular craniotomy was performed unilaterally. The LH was injected at 5.4 mm depth from bregma with either 200 nL of AAV1-hORX.C1V1(t/s).mCherry (> 10^13^ GC/ml) for stimulation experiments, and with 300 nL AAV1-hORX.GCaMP6s (2.5 × 1012 GC/ml) for photometry, 2-photon recordings, as well as controls for the channel-rhodopsin photostimulation stimulation experiments (where GCaMP served as a non-opsin control virus as in previous work (7)). Either 2 optic fibers (0.2mm, Thorlabs) or a GRIN lens (0.6mm, Inscopix) were lowered to the injection locations and cemented in place along with custom made headplate (Protolabs). To specifically target noradrenaline cells for the LC photometry experiments, two 200 nL injections of AAV.CAG.Flex.GCaMP6s.WPRE.SV40 (1.9 x 10^13^ GC/ml, Addgene), were performed in the LC of C57BL/6-Tg(Dbh-icre)1Gsc mice (Jax 4355551, n = 6), unilaterally (0.9 mm lateral and 5.4 posterior from Bregma, at depths of 3.5 and 3.7 mm), where a fibre was implanted and cemented in place. Animals were allowed a minimum of 3 weeks to express the viruses post injection before the experiments started.

### Pupil size measurement

In all experiments pupils were recorded using an infrared camera (FLIR) at a frame rate of 20 Hz. Pupil size was determined using by finding point estimates of 8 points at the edge of the pupil in each frame, using DeepLabCut (27). During blinking or low confidence estimation of points, points were interpolated. A circle was then fitted to the estimated points in Matlab, to determine pupil surface area and radius.

### Optogenetic stimulation

Optogenetic stimulation of orexin neurons was performed using two green lasers (532nm, Laserglow) via an 0.2 mm diameter optic fibre. Light intensity measured at the fibre end was 10 mW and the 5 ms pulses were delivered at different frequencies. During stimulation experiments, animals were kept anesthetized with 2% Isoflurane and delivered a 30 s optogenetic stimulation train at one of the four frequencies (1,5,10 and 20Hz), in a randomised order, every 2-2.5 minutes.

### Pharmacological experiments

Orexin receptor blockade was performed with 100 mg/kg i.p. injections of dual orexin receptor antagonist Almorexant (ALM) dissolved in 10% DMSO vehicle and PBS. Mice were injected with either ALM or vehicle on separate days, 1 hour before anesthetizing and performing optical stimulation. Stimulation was performed as described above.

### Studies of mice without orexin neurons

For complete ablation of orexin neurons, we used mice expressing the human diphtheria toxin (DTR) receptor in orexin cells (orx-DTR mice), DT injection in this mouse model produces complete orexin cell loss while sparing surrounding cell types, as previously described and validated (28). Orexin neurons were deleted in orx-DTR mice with two i.p. injections of 150 ng diphtheria toxin (Sigma D-564) diluted to 1 μg/ml in saline, two days apart. Wild type mice, used as control, were DT injected and analyzed in the same manner. After allowing two weeks to produce complete orexin deletion (28), each mouse underwent three 1-hour long pupil recording sessions, performed on separate days. During recording, the animal was head-fixed to a custom-made post and allowed to run freely on a rotating wheel with a diameter of 20 cm. Running speed was captured using a rotary encoder attached to the wheel and the pupil was recorded at 20 frames per second using a Blackfly FLIR camera. In subsequent analysis, to account for any periods of atonia that can occur after orexin deletion, we analysed pupil size during both running and norunning epochs, using a binary threshold set at >1 cm/s.

### Volumetric 2-photon imaging of single orexin neurons

Imaging was performed as described in our previous work5, using a custom electro-tunable lens equipped resonant/galvanometer scan head two-photon microscope (INSS) and a femtosecond-pulsed mode-locked Ti: sapphire laser (Spectra-physics Mai Tai HP Deepsee 2) at 950 nm through a 20× (0.45 NA, Olympus) air-IR objective at 31 frames/s. Custom Labview software was used to capture 512 by 512 pixel images of neurons through the implanted GRIN lens and a 510/80 nm band-pass emission filter. Six z-planes were imaged with the electro-tunable lens, leading to a volume rate of 5.15 volumes/s. During imaging of orexin neurons in the LH, mice (n = 3) were head-fixed, allowed to run freely on a wheel, and given 5 μL rewards of strawberry milkshake through a spout placed by their mouth. Rewards were delivered in random intervals of 1 - 1.5 minutes, with a total of 50 rewards delivered in a session, using a solenoid valve (NResearch 161K011). The metal reward spout was connected to a custom-made capacitive lick detector. Because pupil diameter in awake mice was too large to monitor in the dark, we elicited mild constriction with a constant weak blue LED light directed at the eye. To simultaneously track pupil size and accurately synchronize signals, a Blackfly FLIR camera frame capture was synced to a single plane capture of the two-photon microscope. Images acquired in imaging sessions were first motion corrected using the TurboReg plug-in for ImageJ software (NIH). Regions of interest (ROIs) were drawn around each cell manually in ImageJ and the mean intensity was extracted from each ROI to get the raw fluorescence of each cell. To correct for neuropil contamination, we also extracted mean intensity of a halo surrounding each cell ROI with a distance of 6 to 12 pixels, but not including other ROIs. Finally, to get the final dF/F signal we subtracted the mean neuropil ROI intensity from the cell ROI intensity. Each z-plane in the volume was inspected to find cells appearing in multiple planes, and dF/F traces belonging to the same cell were averaged together. Each dF/F trace was smoothed with a 3-sample moving average and z-scored. The collected dataset was comprised of dF/F traces from 150 neurons.

### Encoding model

The applied encoding model was based on a previous study (18), and was based on multiple regressions with dF/F trace of each cell as a dependent variable and acquired running, reward and pupil size traces as independent variables (Fig. 3d). Running speed was acquired by a rotary encoder attached to the running wheel. Reward consumption was quantified as the square wave licking signal acquired from the capacitive lick detector convolved with a spline. Spline was picked from a previously used (18) set of splines, picked the one that explained the most variance for most cells in the dataset in subsequent regressions. All predictor traces were z-scored. Having run the regressions, we removed all the cells for which model predicted <5% of total variance. For the remaining cells we re-ran the model with one of the variables removed to estimate their impact on the explained variance. By dividing full model r^2^ with the r^2^ from each of the variable-excluded models, we were able to calculate the contribution of each variable to the variance explained in the full model.

### Fiber photometry

Fiber photometry (Supp. Fig. 2) was performed as in our previous work (7). Photometry signals were detrended by fitting a convex hull around the raw photometry trace such that, moving forwards in time, monotonically decreasing vertices were saved into a template. The template was linearly interpolated to match the length of the photometry trace, and then subtracted from it. Following detrending, all photometry traces were z-scored.

### Coherence analysis

Coherence between photometry and pupil-dilation was calculated via the multitaper method using custom python scripts built upon the NiTime library (https://nipy.org/nitime/). First, photometry and pupil traces were clipped to the same length (approx. 38 minutes) across all mice. Coherence at each frequency was calculated from adaptive weighting of the first 7 tapers in order to target a constant resolution bandwidth. Thereby, coherence was computed with an epoch length of the entire trace, and then averaged across mice.

### Statistical analysis

Unless otherwise specified, all raw data processing and statistical analysis was done in Matlab. The statistical tests and their results are shown in the figure legends, along with n values the tests were based on. P values of <0.05 are indicated with * and values < 0.01 are indicated with **, with all values below the value of 0.05 accepted as significant. All error bars show standard error of the mean (SE).

## ACKNOWLEDGEMENTS

We would like to thank Antoine Adamantidis, Rafael Polania, and Johannes Bohacek for helpful comments on initial drafts. Work was supported by ETH Zurich.

## AUTHOR CONTRIBUTIONS

N.G. and D.B conceptualized and designed the experiments and wrote the manuscript. N.G. performed surgeries, collected data, and carried out data analyses. A.T. and E.B. contributed to the experimental code, performed some of the surgeries and data analysis. D.B. conceived the original hypotheses. All authors contributed to discussion of the results and manuscript. D.R.P. and D.B. raised funding and supervised the study.

---

**Supplementary Figure 1.**
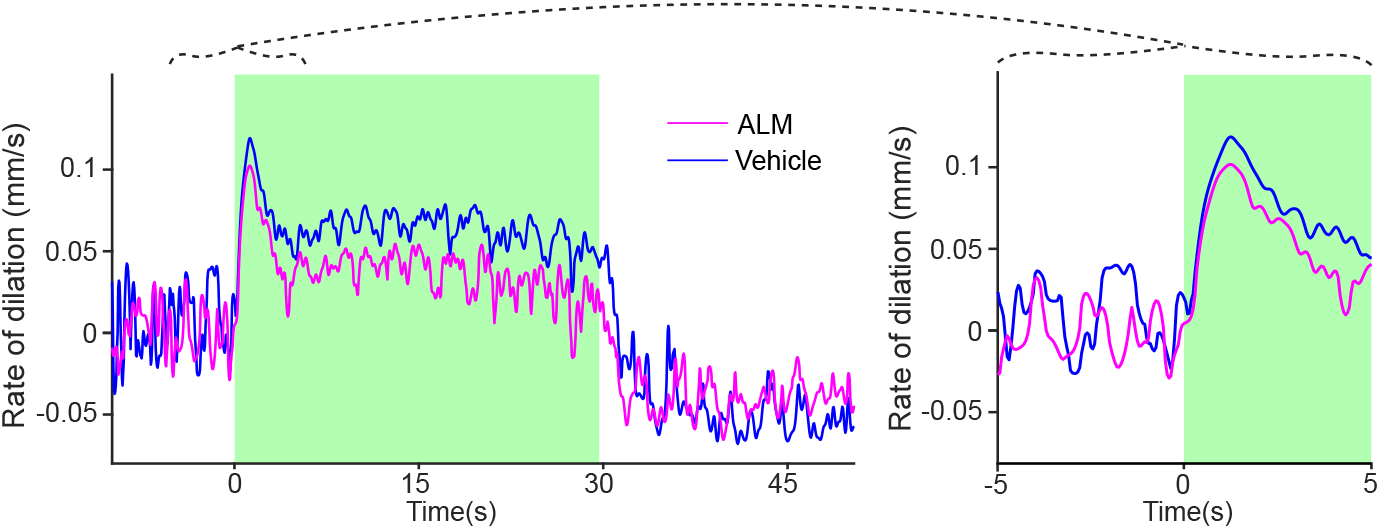
Dilation rates during orexin cell photo-stimulation and orexin receptor blockade. Mean rates of dilation across mice during 20 Hz photo-stimulation of orexin cells in the LH. Shaded green area indicates stimulation duration. Left, dilation rates for Almorexant-injected animals (n = 4 mice, pink) vs. same animals after vehicle injection (blue). Right, zooms of 5 seconds before and after stimulation onset.

**Supplementary Figure 2.**
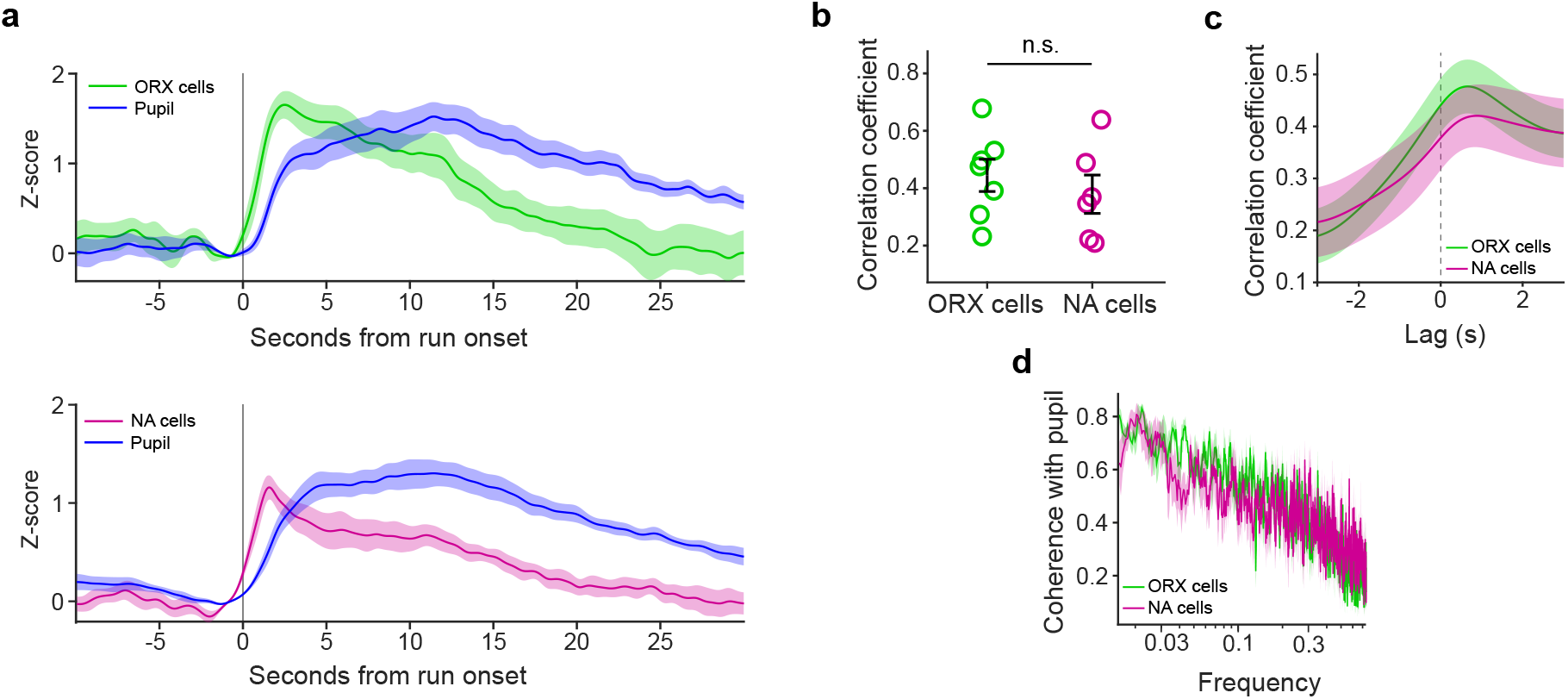
Orexin and LC neurons show similar relationships to pupil size. **a)** Run onset aligned traces of pupil and either orexin cells (ORX, top panel, n = 7 mice) or LC noradrenaline cells (NA, bottom panel, n = 6 mice) photometry signals. **b)** Lag corrected correlation coefficients between pupil size and orexin/LC neuron activity (one-tailed t-test: t = −0.76, p = 0.23). **c)** Cross-correlation plot between pupil and respective neural signal. d, Coherence analysis between pupil and respective neural signals. All error bars show SE.

**Supplementary Figure 3.**
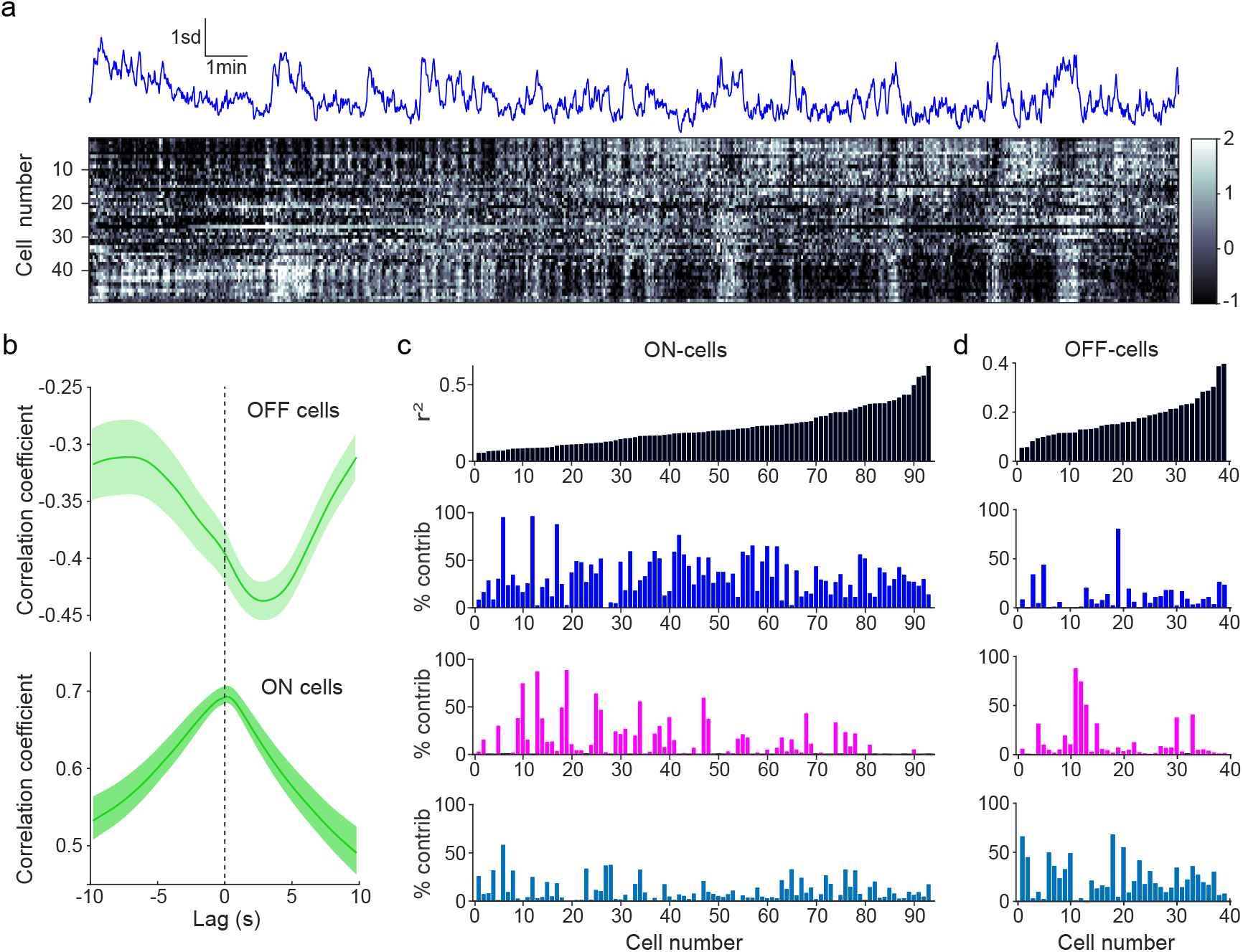
Separate analysis of pupil-ON and pupil-OFF orexin cells. **a)** Top, z-scored pupil trace aligned to (bottom), heatmap of orexin neuron activity arranged by correlation with pupil size from one mouse. **b)** Average of cross-correlations of pupil with cells positively correlated with pupil (ON-cells, bottom, n = 92 cells) or negatively correlated with pupil (OFF-cells, top, n = 40 cells). Shaded areas show 1 standard error. **c)** R-squared values and contributions to variance explained in analysis shown in Fig. 3. Separated for ON-cells (left) and OFF-cells (right).

**Supplementary Figure 4.**
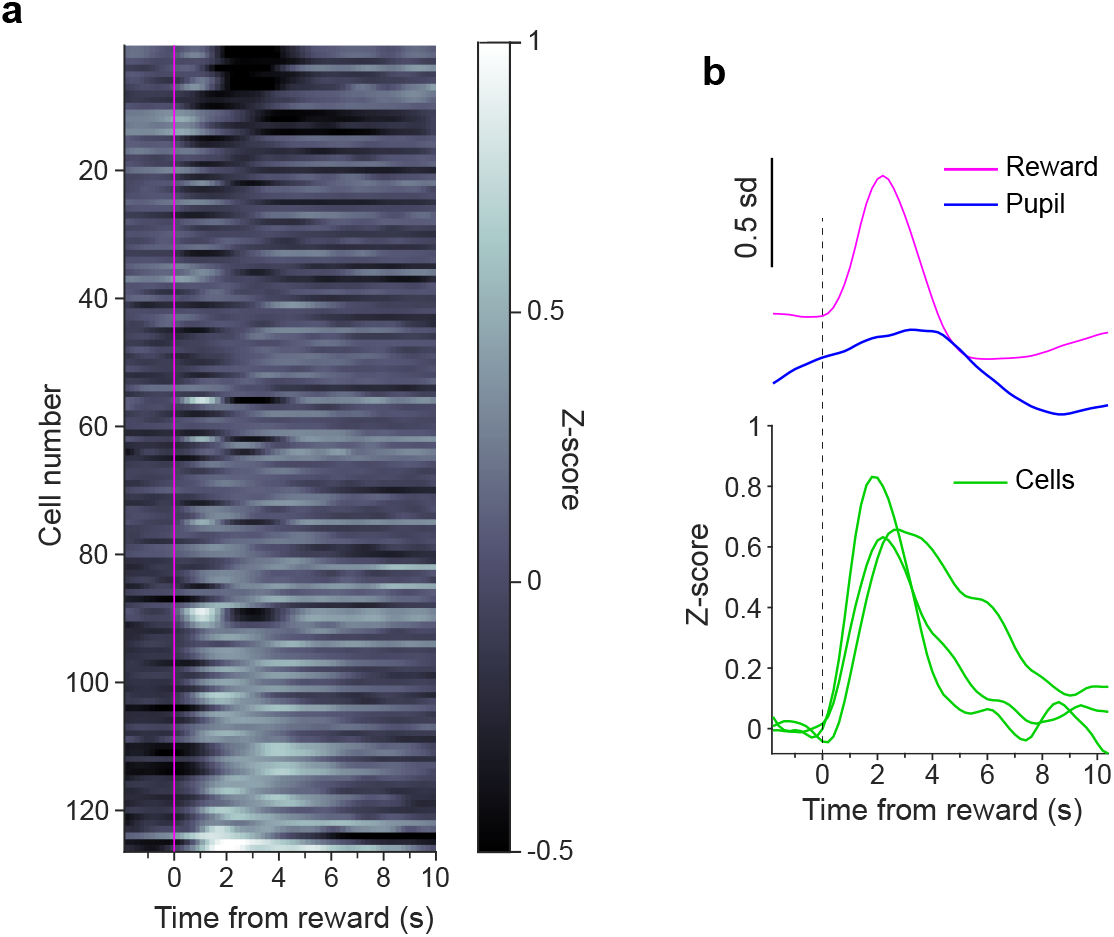
Orexin neurons aligned to reward onset. **a)** Reward onset aligned, trial-averaged and baseline subtracted orexin cell activity for experiment shown in Fig. 3. **b)** Top, example pupil and splined licking trace from one animal. Bottom, reward-onset aligned traces from three orexin cells shown in heatplot in a.

## Notes

### Competing Interest Statement

The authors have declared no competing interest.

